# *In Silico* Analysis of Ixodid Tick Aqauporin-1 Protein as a Candidate Anti-Tick Vaccine Antigen

**DOI:** 10.1101/636985

**Authors:** Christian Ndekezi, Joseph Nkamwesiga, Sylvester Ochwo, Magambo Phillip Kimuda, Frank Norbert Mwiine, Robert Tweyongyere, Wilson Amanyire, Dennis Muhanguzi

## Abstract

Ticks are arthropod vectors of pathogens of both Veterinary and Public health importance. Ticks are largely controlled by acaricide application. However, acaricide efficacy is hampered by high cost, the need for regular application and selection of multi-acaricide resistant tick populations. In light of this, future tick control approaches are poised to rely on integration of rational acaricide application and other methods such as vaccination. To contribute to systematic research-guided efforts to produce anti-tick vaccines, we carried out an *in silico* tick Aquaporin-1 protein (AQP1) analysis to identify unique tick AQP1 peptide motifs that can be used in future peptide anti-tick vaccine development. We used multiple sequence alignment (MSA), motif analysis, homology modeling, and structural analysis to identify unique tick AQP1 peptide motifs. BepiPred, Chou & Fasman-Turn, Karplus & Schulz Flexibility and Parker-Hydrophilicity prediction models were used to asses these motifs’ abilities to induce antibody mediated immune responses. Tick AQP1 (MK334178) protein homology was largely similar to the bovine AQP1 (PDB:1J4N) (23% sequence similarity; Structural superimposition RMS=1.475). The highest similarities were observed in the transmembrane domains while differences were observed in the extra and intra cellular protein loops. Two unique tick AQP1 (MK334178) motifs, M7 (residues 106-125, *p*=5.4e-25) and M8 (residues 85-104, *p*=3.3e-24) were identified. These two motifs are located on the extra-cellular AQP1 domain and showed the highest Parker-Hydrophilicity prediction immunogenic scores of 1.153 and 2.612 respectively. The M7 and M8 motifs are a good starting point for the development of potential peptide-based anti-tick vaccine. Further analyses such as *in vivo* immunization assays are required to validate these findings.

## Background

Ticks are arthropod vectors of pathogens of both Veterinary and Public health concern in tropical and sub-tropical regions of the world [1,2]. In 2007, the average annual cost of tick borne diseases (TBDs) management was estimated to be 4.20 US Dollars per head of cattle [3]. Ticks are associated with both direct and indirect constraints to livestock health and production [1,2]. Their direct constraints to livestock production include, but are not limited to, blood loss (anemia), discomfort, skin-and-hide quality loss, and tick paralysis (Fig 1) [4]. Their indirect constraints to livestock production and health relate to transmission of diseases of veterinary importance including: anaplasmosis (*Anaplasma spp*), babesiosis (*Babesia spp*), theileriosis (*Theileria spp.*), and heartwater (*Cowdria ruminatium*) [1]. In addition, ticks transmit zoonotic pathogens belonging to *Babesia spp*., *Borrelia spp*., *Rickettsia spp*., *Ehrlichia spp*., *Francisella tularensis*, *Coxiella burnetii*, and viruses such as Nairovirus (*Bunyaviridae*) that causes Crimean-Congo hemorrhagic fever (CCHF) and tick-borne encephalitis virus (*Flaviviridae*) [5,6].

**Fig 1.**
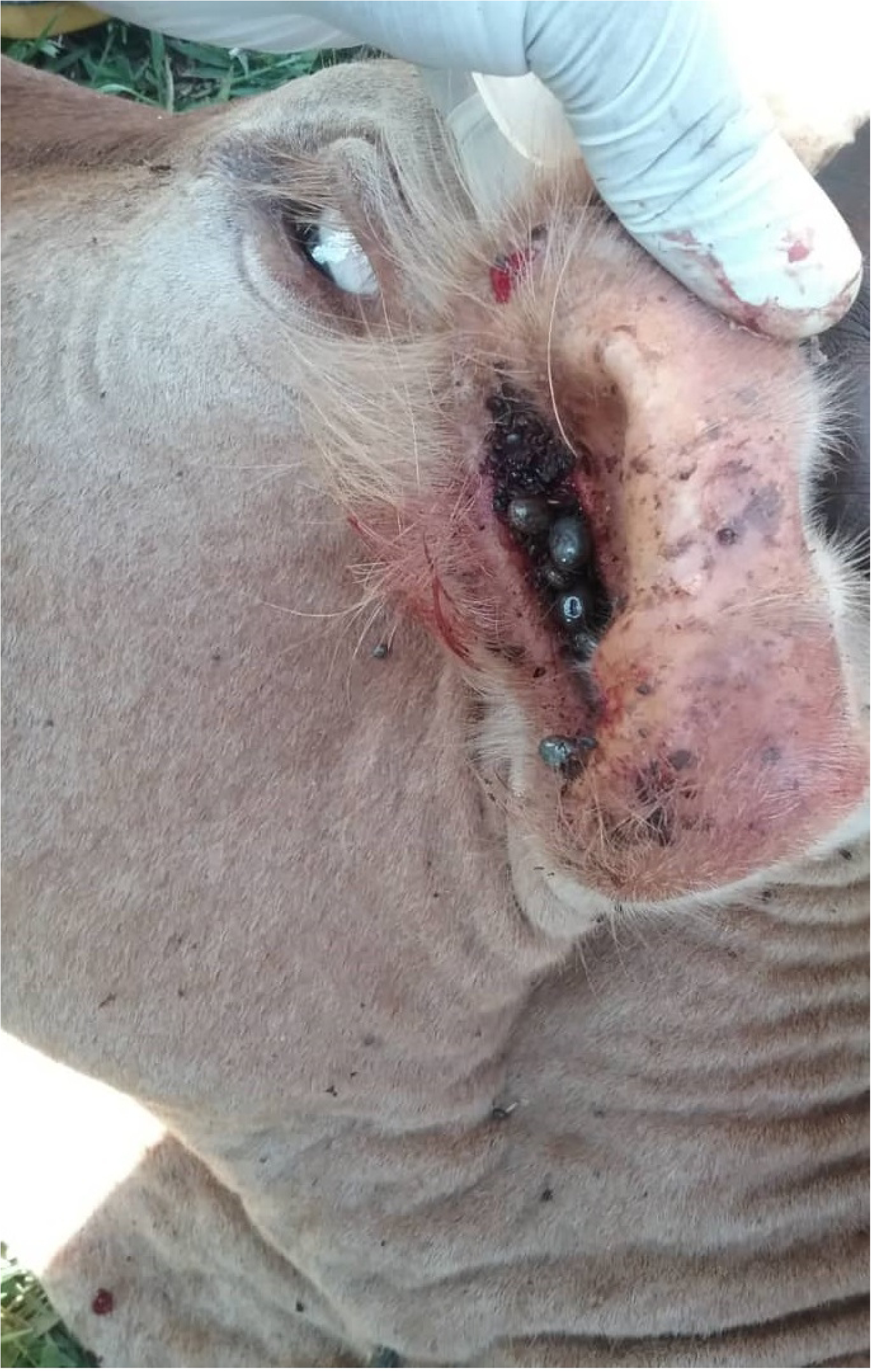
Cattle tick infestation. The image was taken from Serere district Uganda (one of our study districts)

In order to reduce the impact of ticks on livestock health and production as well as withdraw their negative public health effects, acaricides, tick handpicking, animal movement control, and vaccination are often applied to tick control strategies [7]. Acaricides are the most effective control measure of ticks and of tick-borne diseases (TBDs) [8]. However, acaricides’ ever increasing cost in less developed countries, the most affected by ticks and TBDs means; that acaricides cannot be regularly applied for tick control [9]. Moreover, acaricide overuse and misuse are associated with selection of multi-acaricide resistant tick populations [10]; a problem that has been on the rise in Uganda [11]. In light of this, future tick control strategies will have to depend on integration of economically effective acaricide application, vaccination, breeding livestock for tick resistance, and other available tick control methods such as controlled animal movements [9]. To date anti-tick vaccine development has been slow, for example the only BM86 recombinant vaccine against *R. microplus* was developed more than 2 decades ago [12]. There is therefore a need to rekindle systematic research-guided efforts to evaluate crucial tick proteins and biological pathways for vaccine development.

Tick aquaporins belong to the Membrane Intrinsic Protein (MIP) superfamily which are known to play a key role in transport of water and glycerol across the cell membrane. MIP has three subfamilies; Classical aquaporin (cAQP), aquaglyceroporin and S-aquaporin. The major difference between the three subfamilies is in the signature residues around the protein pore. Classical aquaporin allows only the passage of water, while aquaglyceroporin take in glycerol. In ticks, there are two types of aquaporins namely: AQP1 and AQP2 which help in concentrating blood meals by facilitating the excretion of excess water back into the host via the salivary glands [13,14]. From mRNA analysis of female *I. Ricinusi* ticks, it was shown that AQP1 is expressed in tissues such as the gut, rectal sac and most abundantly in the salivary glands where it is involved in water flux, while AQP2 is only expressed in the salivary glands [15]. Additionally, *R. microplus* AQP1 recombinant antigens showed 75% and 68% efficacy against adult female ticks in two trials involving groups of one-year old calves [16]. Lastly, vaccination of rabbits with a conserved region of *I. ricinus* larval APQ1 in rabbits has previously been shown to provide 80% tick reduction highlighting the potential of using AQP1 as a candidate antigen for anti-tick vaccine development [17].

The development of an AQP1 derived anti-tick vaccine remains a challenge mostly due to the high sequence and structural similarity among tick and host (humans and Bovine) AQP1 orthologs [16]. However, this can be overcome by identifying tick specific immunogenic regions within the AQP1 protein that will allow for a more selective immunogenic response. Furthermore, tick species isolated from different tick endemic regions are required to assess the conservation of these AQP1 immunogenic regions. In the study herein, we focused on analysing tick AQP1 from *Ixodid* ticks isolated from three different agro-ecological zones in Uganda and bovine AQP1 proteins derived from Protein Data Bank (PDB) in order to identify unique tick AQP1-peptide motifs that can be used in future anti-tick vaccine development pipelines.

## Methods

### Study design

This study involved identification and confirmation of representative tick species from ticks isolated from the four main cattle keeping districts of Uganda. After which, the AQP1 gene was amplified using PCR and sequenced using Sanger method [18]. To identify unique tick AQP1 peptide motifs, we used multiple sequence alignment, motif analysis, homology modeling and structural analysis. Humoral (B-cell) epitope prediction models, BepiPred, Chou & Fasman-Turn, Karplus & Schulz Flexibility and Parker-Hydrophilicity prediction models were used to asses these motifs’ abilities to induce antibody mediated immune responses (Fig 2).

**Fig 2.**
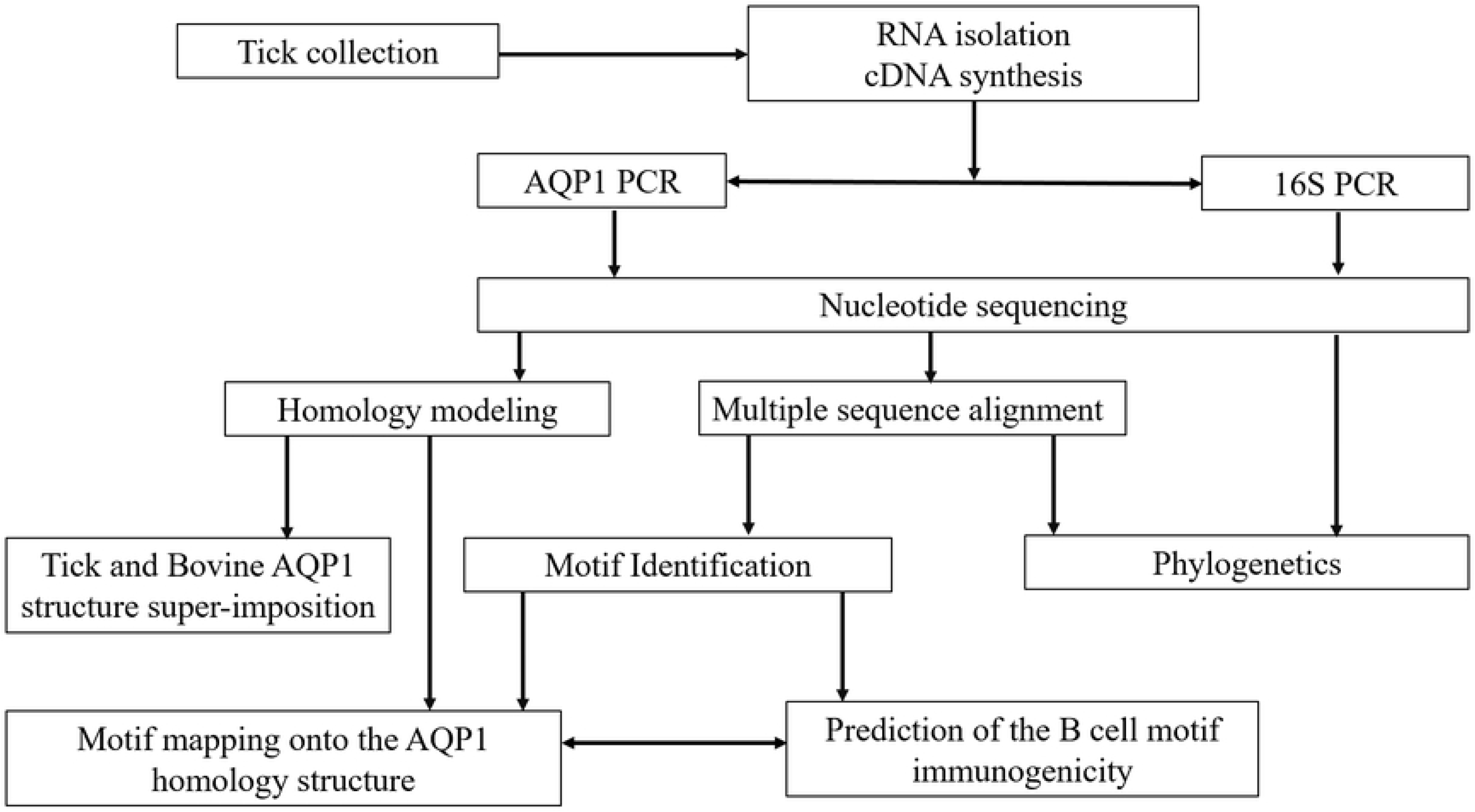
Bioinformatics analysis of unique tick AQP1 motif pipeline. Unique tick APQ1 motifs were detected in a series of steps. First, ticks were collected and identified from 4 district of Uganda. Second the AQP1 gene was amplified in each tick species. The amplified gene from this study and those from the GenBank were further compared. In depth analysis of tick and bovine AQP1 amino acids was done to assess whether there is any difference between these protein orthologs. AQP1 homology modeling and structural mapping were then done to locate each of the predicted peptide motifs on the modeled protein

### Study area description

Ticks used in this study were collected from four districts of Uganda namely; Serere, Kotido, Mbarara and Kiruhura district (Fig 3). These districts were selected basis on the fact that they are in a tick endemic cattle corridor. [19]. These districts are from 3 different regions of Uganda, namely north-eastern Uganda (Kotido district), in south-eastern Uganda (Serere district) and south-western Uganda (Kiruhura and Mbarara districts). These regions represent the three main cattle keeping agroecological zones of Uganda.

**Fig 3.**
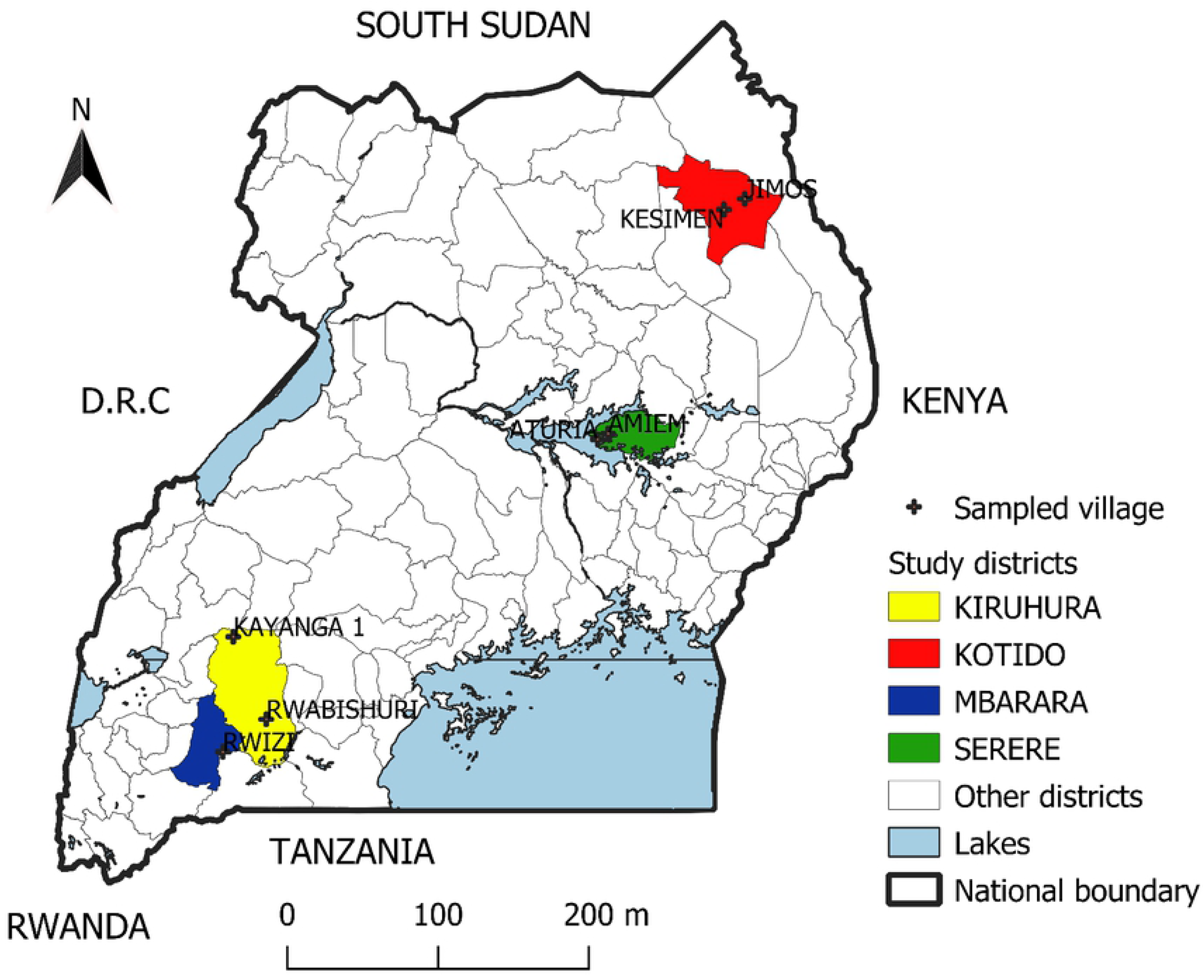
Map of Uganda showing the study districts and villages where ticks were collected. Ticks were collected from 4 districts in the cattle corridor that harbours over 80% of Uganda’s cattle population. The districts include Kiruhura, Mbarara, Serere and Kotido.

### Tick collection and preservation

Cattle in each village were gathered at central cattle holding ground. Cattle were physically restrained before half body tick collection. Ticks were collected from each of the five predilection sites namely; i) the inner and outer fore-legs, ii) hind-legs and abdomen, iii) tail and anal area, iv) head and neck, v) lateral area and dorsal area shoulders to tail base and ears. Collected ticks were put in different customised tick collection containers covered with gauze and transferred to Makerere University, College of Veterinary Medicine, molecular biology laboratory for morphological and molecular identification to species level.

### Morphological and Taxonomic tick identification

Each tick was morphologically identified to species level using morphological keys as previously described [20,21].

### RNA extraction and cDNA synthesis

Total RNA from homogenized tick samples was extracted using GE healthcare RNA extraction kit (Chicago, USA). Copy DNA (cDNA) was synthesized using a GoScript™ Reverse Transcriptase (Promega, 2800 Woods Hollow Road Madison, WI 53711-5399 USA). Briefly, 1µl of RNA, 5µl of poly T primers and 14µl of nuclease free water was preheated at 70°C then 25°C for 10 min each. The mixture was placed on ice prior to further extraction steps. The enzyme mix containing 1× first strand buffer, 4µl of MgCl_2_, 2mM dNTP, 0.51µl of Superase inhibitor, 1µl of superscript II, 5µl of the RNA mix, and RNAse free water. The mixture was then incubated in one cycle of 10min at 25°C, 45min at 37°C, 45min at 42°C, followed by 15min at 70°C in a thermocycler. The cDNA was used immediately or stored at −20°C until it was needed for further analysis.

### PCR amplification of 12S and AQP1 genes

Tick species were confirmed by PCR and sequencing of 12S ribosomal RNA (12S rRNA) as previously described [22]. The 12S PCR was carried out using a single pair of primers (12S-F-AAACTAGGATTAGATACCCT) and (12SR-AATGAGAGCGACGGGCGATGT) that amplify a 300bp fragment of the gene. Briefly, PCR reactions were prepared in 50µl mix containing 1.56U Taq DNA polymerase (New England Biolabs, USA) 1× standard buffer, 0.25mM of each dNTP, 0.25 mM each of forward and reverse primers, 1.25mM MgCl_2_, PCR grade water and 5µl of the cDNA template. The gene was amplified in a S1000 ™ Thermal Cycler (BIO RAD, California, United States) with an initial denaturation of 94°C for 5min followed by 30 cycles of 94°C for 30sec, 52°C for 45sec, 72°C for 45sec and a final extension of 72°C for 7min. We designed a single pair of primers (AQP1-1F-5’GCGTGAAGATCAAGAACGCC3’ and AQP1-1R 5’GCCAATTGGAATCGAGGTGC3’) to amplify a 842bp fragment spanning the entire AQP1 gene. The AQP1 PCR reactions were performed in a S1000 ™ Thermal Cycler (BIO RAD, California, United States) in 50 µl final volume containing 1X standard buffer, 0.25mM of each dNTP, 0.25 mM each of forward and reverse primers, 1.56 U *Taq* DNA polymerase, 1.25mM MgCl_2_, 32.18µl of PCR grade water and 5µl of the cDNA template. The thermocycler was for 95°C for 5min, followed by 30cycles of 95°C for 30sec, 50°C for 30sec and 72°C for 1min, and a final extension step at 72°C for 10min.

### Aquaporin-1, and 12S gene amplicon analysis and sequencing

All PCR amplicons were resolved on 2% Agarose gels. The amplicons were sized against a 50bp DNA molecular ladder (Bioline). PCR products were purified using QIAquick^®^ PCR purification kit (Qiagen, USA) according to manufacturer’s instructions prior to sequencing. The PCR amplicons were commercially Sanger sequenced by Inqaba Biotechnical Industries (Pty) Ltd (Pretoria, South Africa).

### Sequence identification, multiple sequence alignment and phylogenetics

BLASTn search was performed on both AQP1 and 12S gene sequences using NCBI against the non-redundant nucleotide DataBank. AQP1 gene sequences were translated into 6 open reading frames using Expassy translate tool [23]. BLASTp was used to search for similar aquaporin AQP1 amino acid sequences against the protein DataBank using BLOSUM62 matrix.

Aquaporin 1 amino acid and 12S nucleotide sequences from this study and respective similar sequences retrieved from the GenBank were aligned together in MUSCLE using default settings [24]. Phylogenetic analyses were performed using the maximum likelihood method with 1000 bootstrap in MEGA software version X [25]. Pearson correlation matrix of aquaporin-1 was done using R studio using Sequir, ggplot2 and reshape2 packages.

### Peptides motif modeling

Bovine and tick aquaporin-1 fasta amino acid sequences were submitted to Multiple Em for Motif Elicitation (MEME) Version 5.0.5 motif discovery tool using default settings (http://meme-suite.org/tools/meme) [26]. The MEME was set to pick at most 50 motifs which appear in at least 2 sequences. Motif heatmap was generated using an in-house python scrip. Tick AQP1 motifs were analysed for their conservation among different tick species using WebLog [27].

### Aquaporin-1 protein homology modeling

The 3D structure of AQP1 was modeled using PRotein Interactive MOdeling (PRIMO) pipeline (https://primo.rubi.ru.ac.za) (using default settings) [28]. Structures with identity percent higher than 30% were selected as suitable template model structures. 1LDF (Method: X-RAY diffraction, Resolution: 2.1 Å, R-Value Free: 0.243, R-Value Work: 0.230) and 1FX8 (Method: X-RAY diffraction, Resolution: 2.2 Å, R-Value Free: 0.223, R-Value Work: 0.197) were chosen as template structure models. Four models were produced using the default settings. These model structures were evaluated in Protein Structure Analysis (ProSA-web) and Ramachandran plot (version 2.0) [29–31]. The modeled motifs were then mapped onto the AQP1 homology structure using PyMol, Version 1.7.4 [32]. Structural super-imposition of the Model AQP1 protein and bovine AQP1 protein (PDB 1J4N, Method: X-RAY diffraction, Resolution: 2.2 Å, R-Value Free: 0.308, R-Value Work: 0.266) were done using the same PyMol software.

### *In silico* B cell binding peptides prediction

Aquaporin-1 motifs were subjected to a semi-empirical method which uses physiochemical properties, hydrophobicity and secondary structures of amino acids residues and their frequencies of occurrence to predict their immunogenicity abilities [33–35]. AQP1 motifs were pasted onto the analysis window of IEDB analysis resources (http://tools.iedb.org/main/). B cell antigenic determinants were evaluated using different methods namely; Bepipred linear epitope prediction, Chou & Fasman Beta-turn prediction, Karplus & schulz and parker Hydrophicity prediction.

#### Ethical review

This study received institutional review board clearance from the Makerere University, School of Biosecurity, Biotechnology and Laboratory Sciences Research and Ethics Committee (SBLS/REC/16/136) and Uganda National Council for Science and Technology (A 513). Informed consent was obtained from cattle owners before their cattle were enrolled into the study. All efforts were made to minimize animal stress during cattle restraint and tick collection. These activities were conducted by qualified registered veterinarians.

## Results

The amino acid sequence numbering used in this section is based on tick AQP1 (Accession: MK334178).

### Tick species confirmation basing on the 12S rRNA

Six main tick species were morphologically identified and confirmed by sequencing the 12S rRNA gene (Fig 4). (Morphological results not presented) *Rhipicephalus turanicus*, *Rhipicephalus muhsamae* and *Amblyomma lepidum* from Kotido district, *Rhipicephalus appendiculatus* from Mbarara, Serere and Kiruhura districts. *Rhipicephalus microplus* and *Amblyomma Variegatum* from Serere district. *R. turanicus* (MK332384), and *A. lepidum* (MK332383) in this study showed very high sequence divergence compared to those in the GenBank.

**Fig 4.**
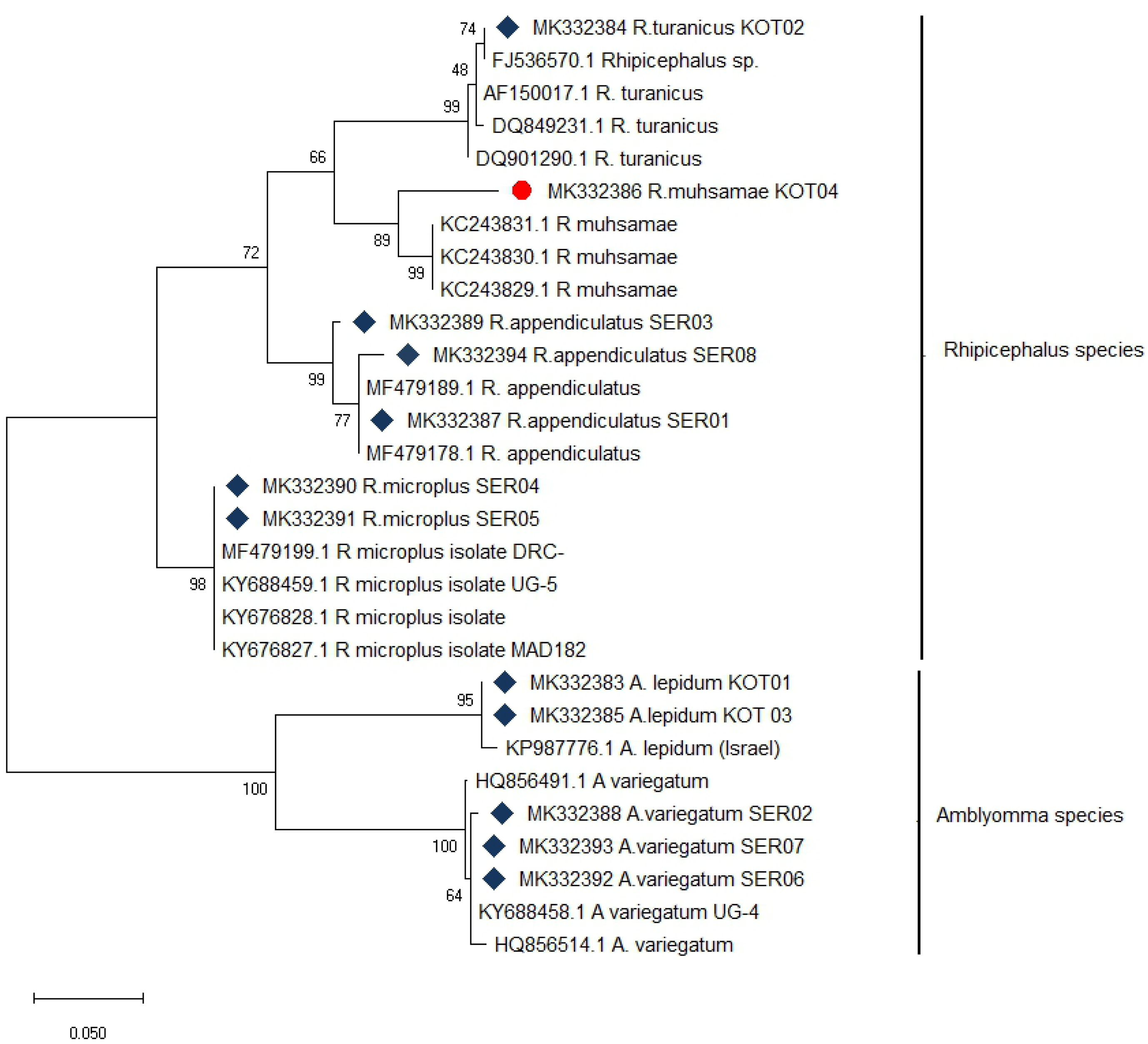
Maximum likelihood phylogenetic tree of tick species based on 12S rDNA gene. Six main tick species were confirmed from different sampling sites (districts). Sequences from this study are identified with 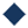 and 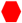 indicates a new tick species identified for the first time in Uganda (this study). The phylogenetic tree was constructed using MEGA X software with the maximum likelihood model with 1000 bootstraps. The scale indicates the rate of nucleotide substitution per node.

### Diversity of Aquaporin-1 protein in different tick species and other orthologs

The amino acid sequences translated from tick AQP1 gene sequences from this study (accession numbers MK334175-MK334178) showed high similarity and with those obtained from the GenBank, with amino substitution rates ranging from 0.025-0.80. Amino acid substitution rate close to 0.00 indicate sequence similarities. Although a high AQP1 similarity was observed among different tick species, a significant amino acid divergence was observed in bovine and human AQP1 sequences when compared with tick AQP1. The evolutionary divergence between ticks and human or bovine AQP1 sequences ranged from 1.110 to 1.270 (calculated using MEGAX). Phylogenetic analysis further confirmed this divergence where tick AQP1 sequences grouped in a distinct clade while human and bovine AQP1 clustered together in a separate clade (Fig 5).

**Fig 5.**
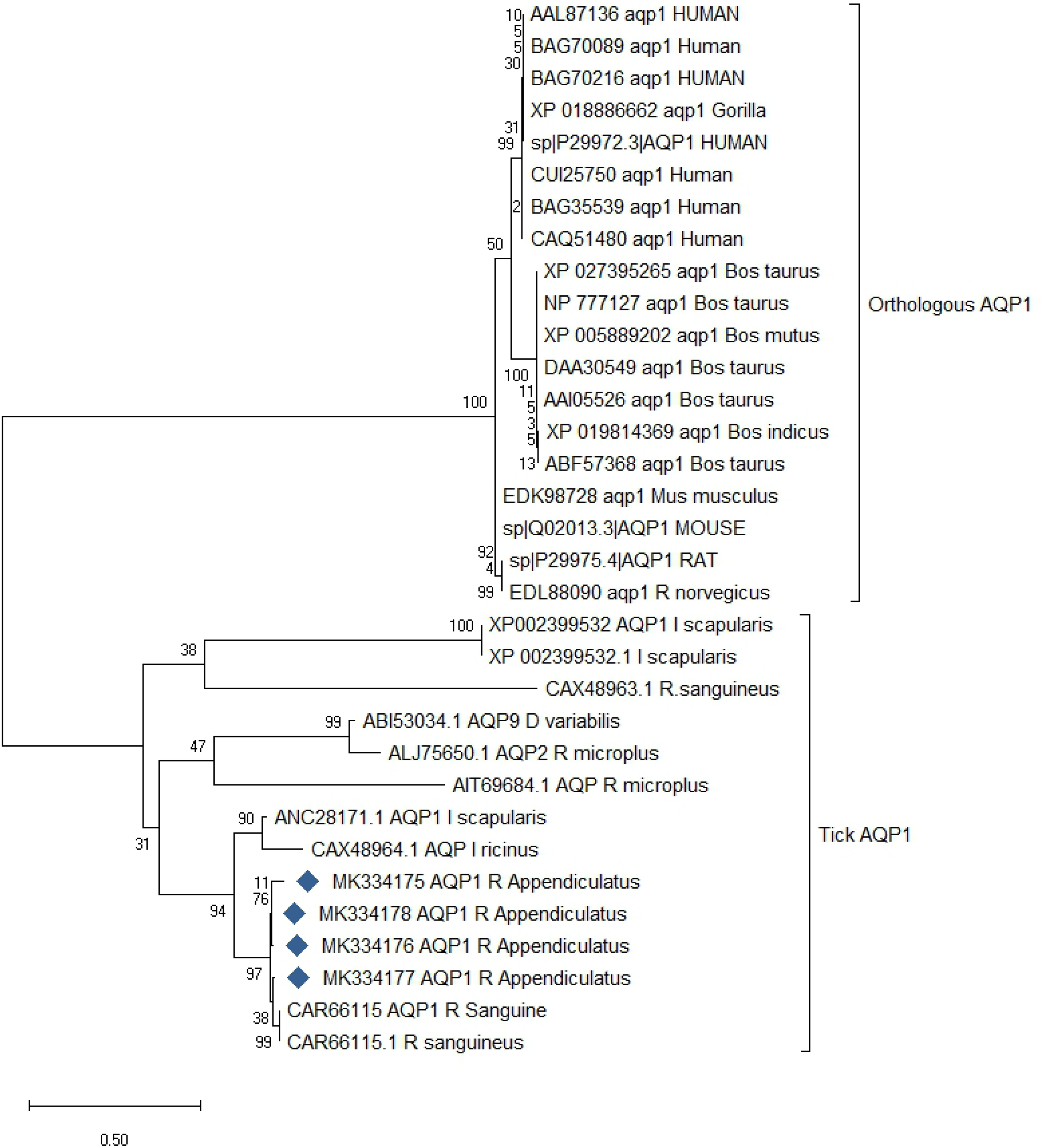
Maximum likelihood phylogenetic tree of AQP1 amino acid sequences from various tick species and other organisms. Ticks and orthologous AQP1 protein sequences clustered in two distinct clades. Tick AQP1 amino acid sequences from this study are identified with 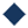. The analysis involved 33 amino acid sequences, which constituted a total of 278 positions in the final dataset. The phylogenetic tree was constructed using MEGA X software with the maximum likelihood model with 1000 bootstraps. The scale indicates the rate of amino acid substitution per node.

The amino acid sequences were additionally analysed using Pearson correlation matrix that revealed that tick AQP1 had a within positive correlation and a negative correlation between bovine and human AQP1 amino acid sequences ranging −0.6 to −0.4 (Fig 6). These results imply that much as AQP1 is conserved among different species, some differences exist between tick AQP1 and bovine or human AQP1 proteins at the sequence level.

**Fig 6.**
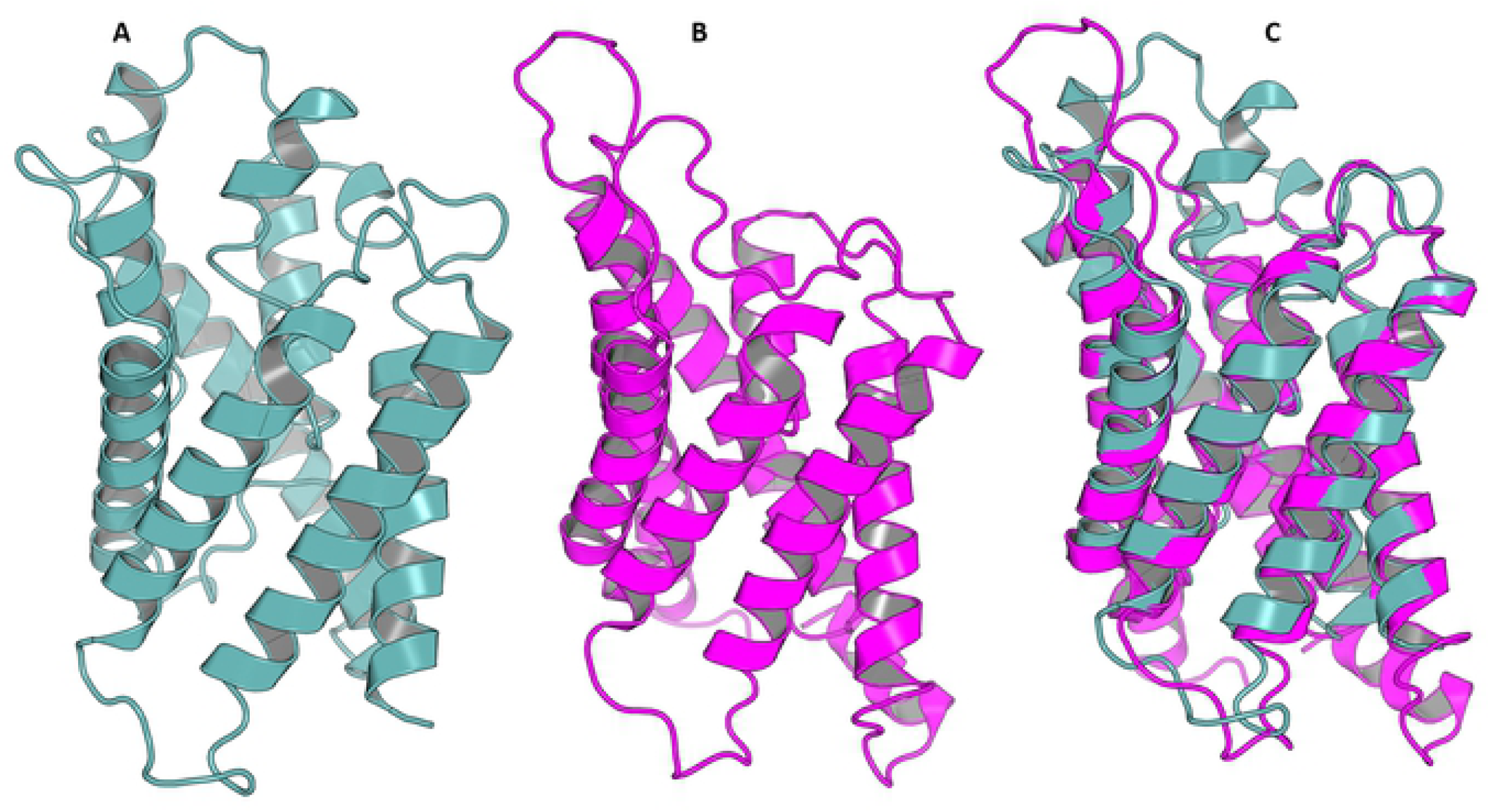
Pearson correlation matrix of aquaporin-1 amino acid sequences. Analysis by Pearson correlation showed that Tick APQ1 has a negative correlation with bovine and human AQP1, highlighting that tick AQP1 has some regions of difference when compared to bovine and human AQP1. A blue color range denotes a negative correlation whereas red indicates a positive correlation. The function scale fill gradient2 was used with the argument limit, c (−1,1).

### Tick and bovine peptide motif modeling

Multiple sequence alignment (MSA) analysis of the tick AQP1 and its bovine ortholog showed some regions of high sequence conservation. The Asparagine-Proline-Alanine (NPA) motif (NPA1: residues 49-51, NPA2: residues 180-181) was conserved in all amino acid sequence orthologs (Fig 7). The NPA is a highly conserved hydrophobic motif that forms part of the pore for each AQP1 monomer [36]. In addition, tick AQP1 motifs contained an aspartic acid (D) residue after the second NPA motif, a signature for aquaglyceroporins (Fig 7).

**Fig 7.**
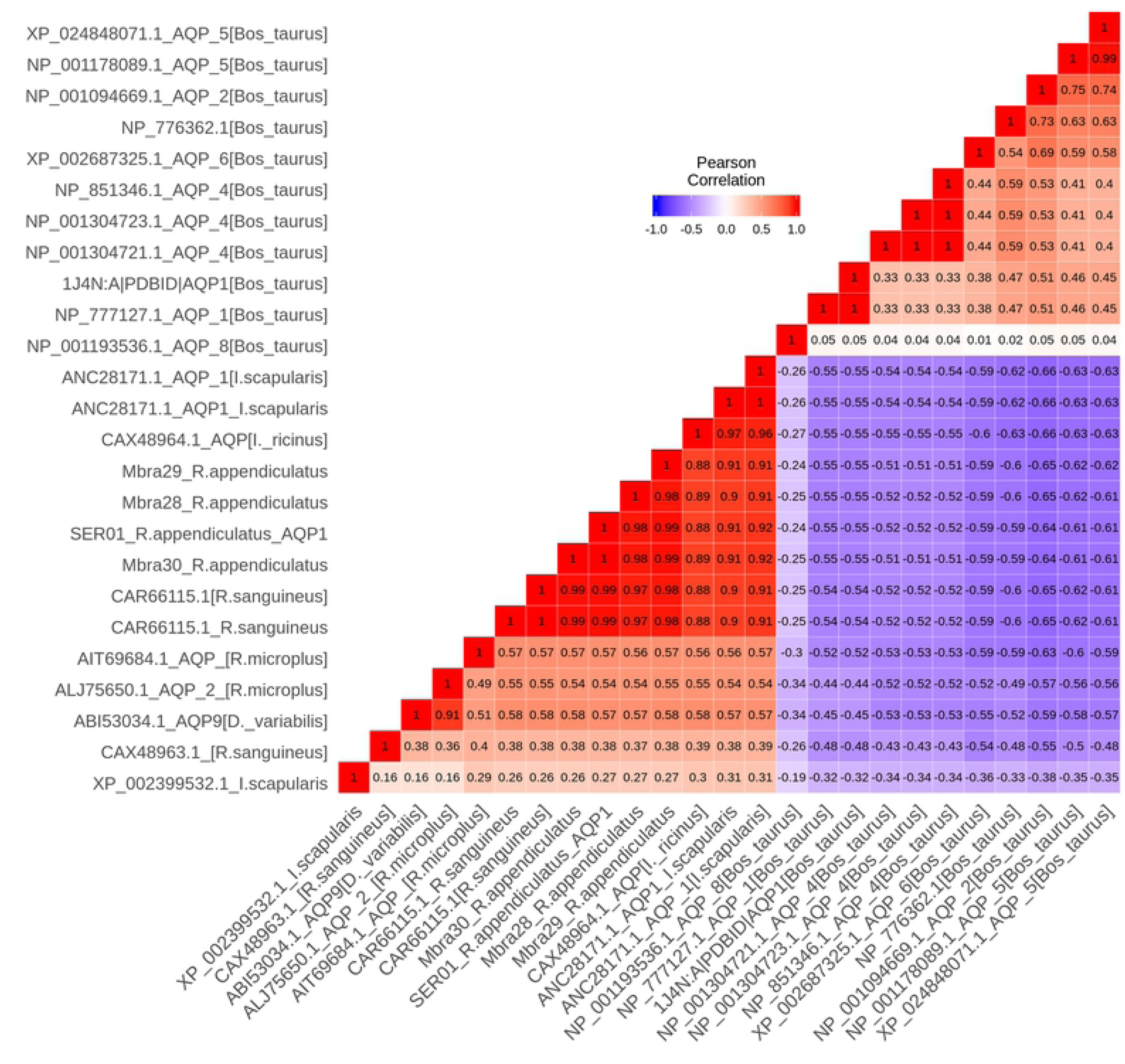
Multiple sequence alignment of tick and bovine aquaporin-1 amino acids. MSA analysis revealed conserved regions within tick AQP1 amino acid sequences absent in bovine AQP1 amino acid sequences. Tick AQP1 has an Aspartic acid residue (D) after the second NPA motif. The MSA was completed using MUSLE. The AQP1 amino acids with the same physiochemical properties were highlighted in same colour code using Clustal.

Motif analysis using MEME revealed six AQP1 motifs located on the extra and intra cellular domains of the homologous tick AQP1 that were unique to ticks (Fig 8). These motifs included; M3 (residues 2-11), M7 (residues 12-31), M8 (residues 85-104), M9 (residues 106-125), M10 (residues 149-168), and M16 (residues 193-212) (Fig 8). All the 6 motifs showed high amino acid sequence conservation across different tick species with E values ranging from 1.6e-108 to 7.6e-013. The position *p* value of the consensus tick AQP1 sequence (MK334178) further confirmed this conservation level across all the six motifs. This *p* value indicates the probability of a single motif appearing in the observed consensus sequence. Position *p* values less than 0.0001 were considered significant.

**Fig 8.**
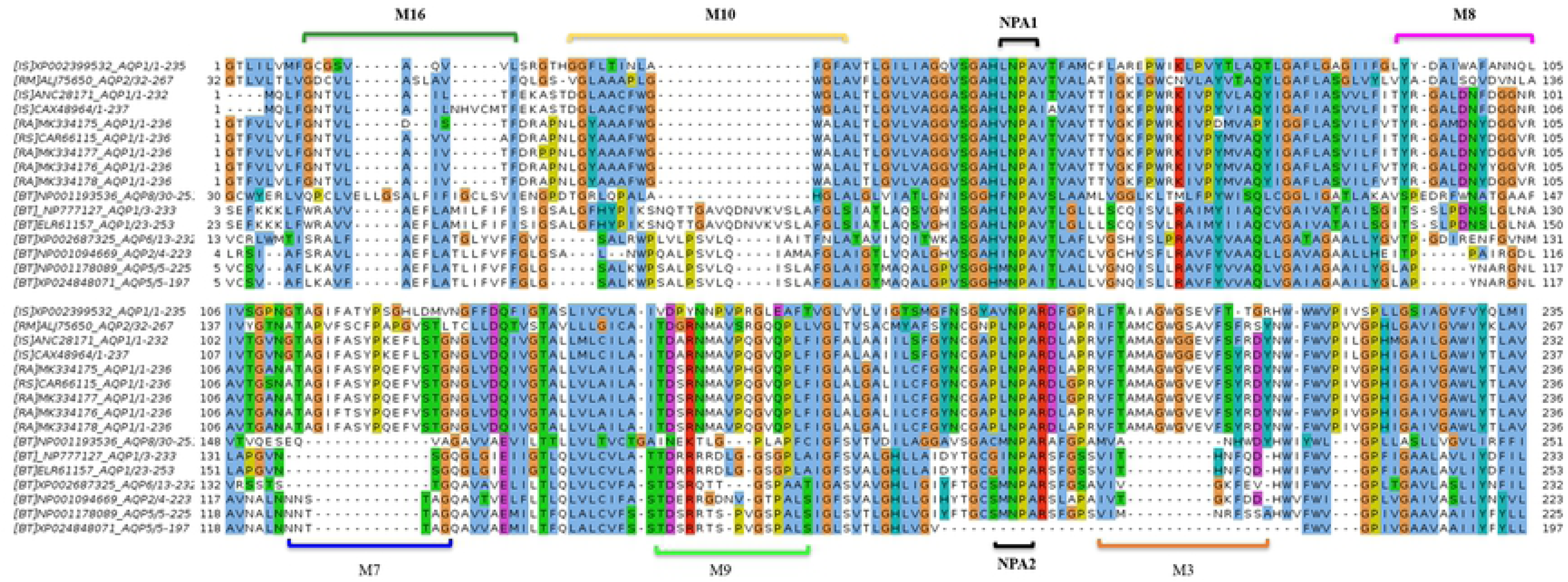
Heat map of tick and bovine Aquaporin-1 peptide motifs. MEME analysis revealed unique tick motifs (M3, M7, M8, M9, M10, M16) denoted by red diamonds. The analysis further showed a number of conserved motifs in both ticks and bovine AQP1 amino acid sequences. The heat map was constructed using an in-house python script.

The conservation level in particular motifs was biased to amino acids with the same physiochemical properties. For instance, motifs M3, M9, M10 consisted of a high number of conserved hydrophobic amino acids (particular in residues 16-20, 9-20 and residues 11-19 respectively) which are colored in black (Fig 9). The most highly conserved motif was M16 with over 70% of its residues being hydrophobic with most of the motif located in the transmembrane domain. Motif M7 contained a high number of substitutions and all these alterations were observed in one tick species (*I. sanguine* sequences XP002399532) with a *p* value of 5.4e-25. Motifs M7 and M8 contained mostly hydrophilic residues (Fig 9).

**Fig 9.**
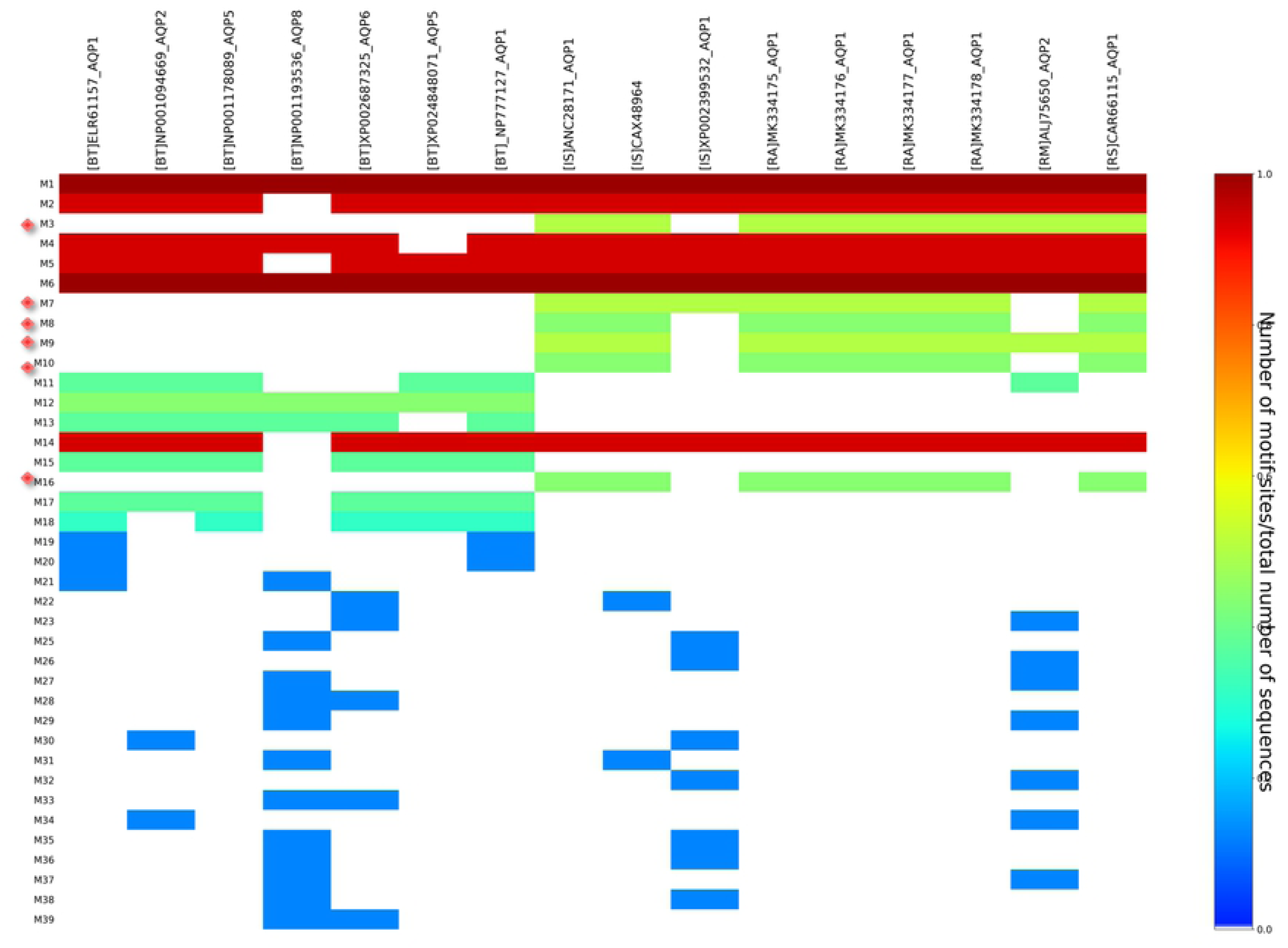
Amino acid sequences of different motifs and their log-sequence. All the motifs were on average 20 amino acids long except motif M16, with 10 amino acids.

### Tick Aquaporin-1 homology modeling and structure analysis

A homology model for tick AQP1 was calculated using PRotein Interactive MOdeling (PRIMO) using *Escherichia coli* (strain K12) AQP (PDB: 1FX8 and 1LDF) as templates. A total of four different models (model01, model02, model03 and model04) were calculated Table 1. Homology model04 (Fig 10-A and 10-B) that had the lowest z-DOPE (normalised Discrete Optimized Protein Energy) score of −0.580 (Table 1) and a ProSA z-score of −3.72 was selected for use in further analyses (Fig 10-C). A z-DOPE score close to −1 indicates that the protein model is close to the native structure [37]. Structure super-imposition of the model proteins and the templates further confirm that model04 is more similar to templates protein (1FX8 and 1LDF) with a Root Mean Square Deviation (RMSD) of 0.34 and 0.36 respectively. A Ramachandran plot score analysis of these models further revealed that model04 had 170 (89.9%) of its residues in the most favoured regions, 16 (8.5%) residues in the additional allowed region and 3 (1.6%) residues in generously allowed region. (Fig 10-D). The NPA1, NPA2 and ar/R filter are shown in Fig 10-A while Fig 10-B shows residues of the amino acid residues of the ar/R filter

**Fig 10.**
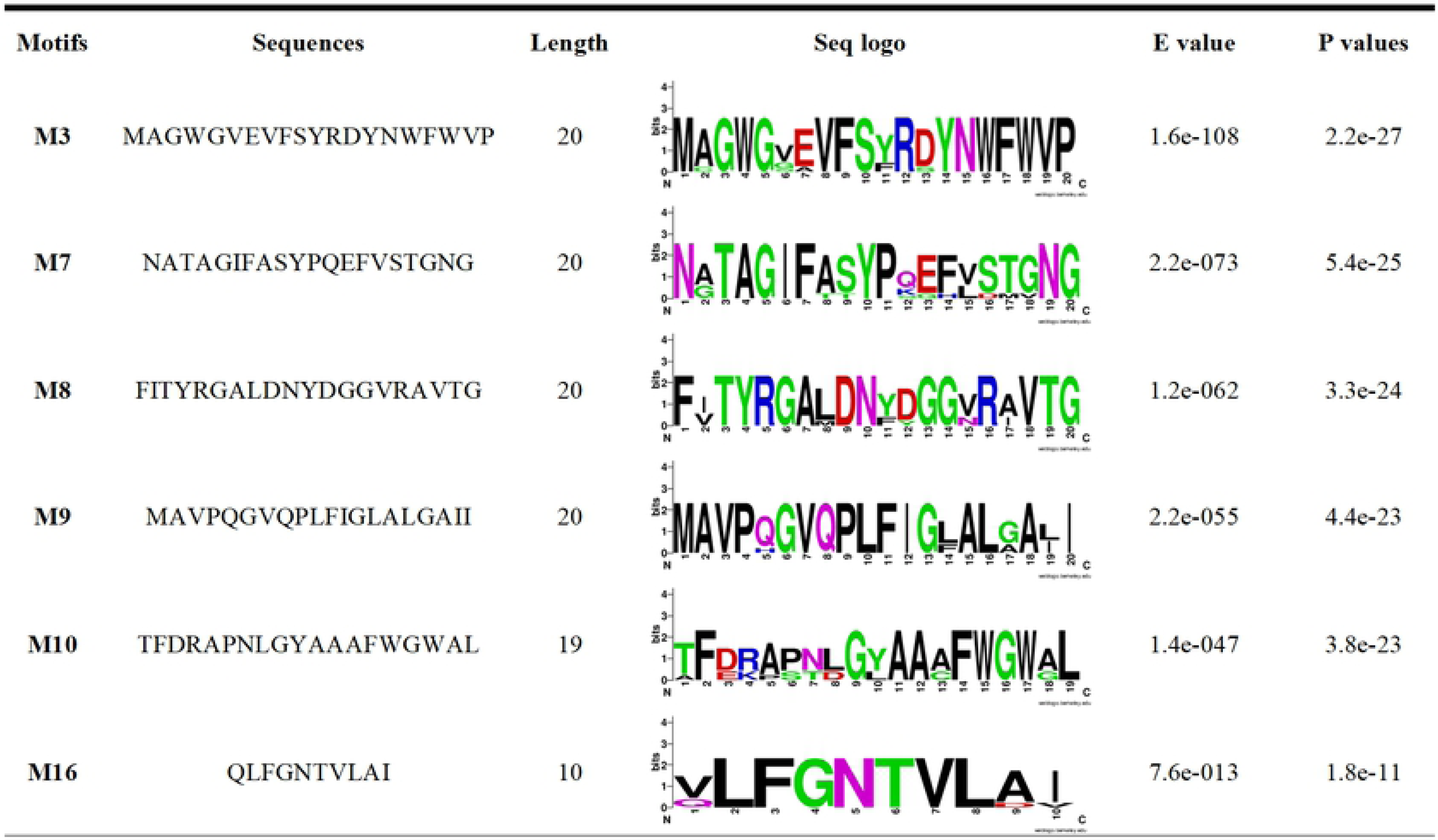
A cartoon showing aquaporin-1 3D structure homology prediction and structure quality analysis. **A.** Monomeric lateral 3D view of the predicted tick aquaporin-1 showing the position of the two Asparagine-Proline-Alanine (NPA) motifs and the Aromatic-arginine (ar/R) filter. **B.** Surface view of tick aquaporin-1 protein molecule showing the amino acid sequences that constitute the ar/R filter. **C.** A ProSA z-DOPE score plot showing the modeled tickAQP1 protein quality (−3.72), **D.** A Ramachandran plot showing the AQP1 residues in the most favoured regions denoted by the red colour. Residues in generously allowed region are indicated with ~a, ~b, ~l, and ~p. Those in additional allowed regions are indicated with a, b, l and p while those in the most favored regions are indicated with A, B, and L. The protein was modeled using PRIMO server using 1FX8 and 1LDF proteins as template structures.

**Table 1.** Comparison of Ticks AQP1homology models and templates AQP1 structure.

| Model name | Dope Z-Score | RMSD |  |
| --- | --- | --- | --- |
|  |  | 1FX8 | 1LDF |
| Model04 | -0.580 | 0.34 | 0.36 |
| Model03 | -0.511 | 0.36 | 0.40 |
| Model02 | -0.370 | 0.44 | 0.43 |
| Model01 | -0.366 | 0.38 | 0.41 |

The homology model with a DOPE Z-score close to -1 indicate homology of good quality. Structure alignment with RMSD close to zero indicate the two protein structures are structural similar

The six motifs were then mapped onto the modeled AQP1 protein structure (model04) to determine their location on the protein 3D structure. Motifs M9, M10, and M16 were localised on the transmembrane protein domain, while motifs M3, M7 and M8 on the extracellular protein domain (Fig 11). The structural superimposition of the tick AQP1 homology model04 (Fig 12-A) and bovine AQP1 (PDB: 1J4N) (Fig 12-B) showed that the proteins were structurally similar with a Root Mean Square Deviation (RMSD) value of 1.475. The structural similarities were mainly observed in the transmembrane domain while differences were observed in the loops located on the extracellular domain of the protein (Fig 12-C).

**Fig 11.**
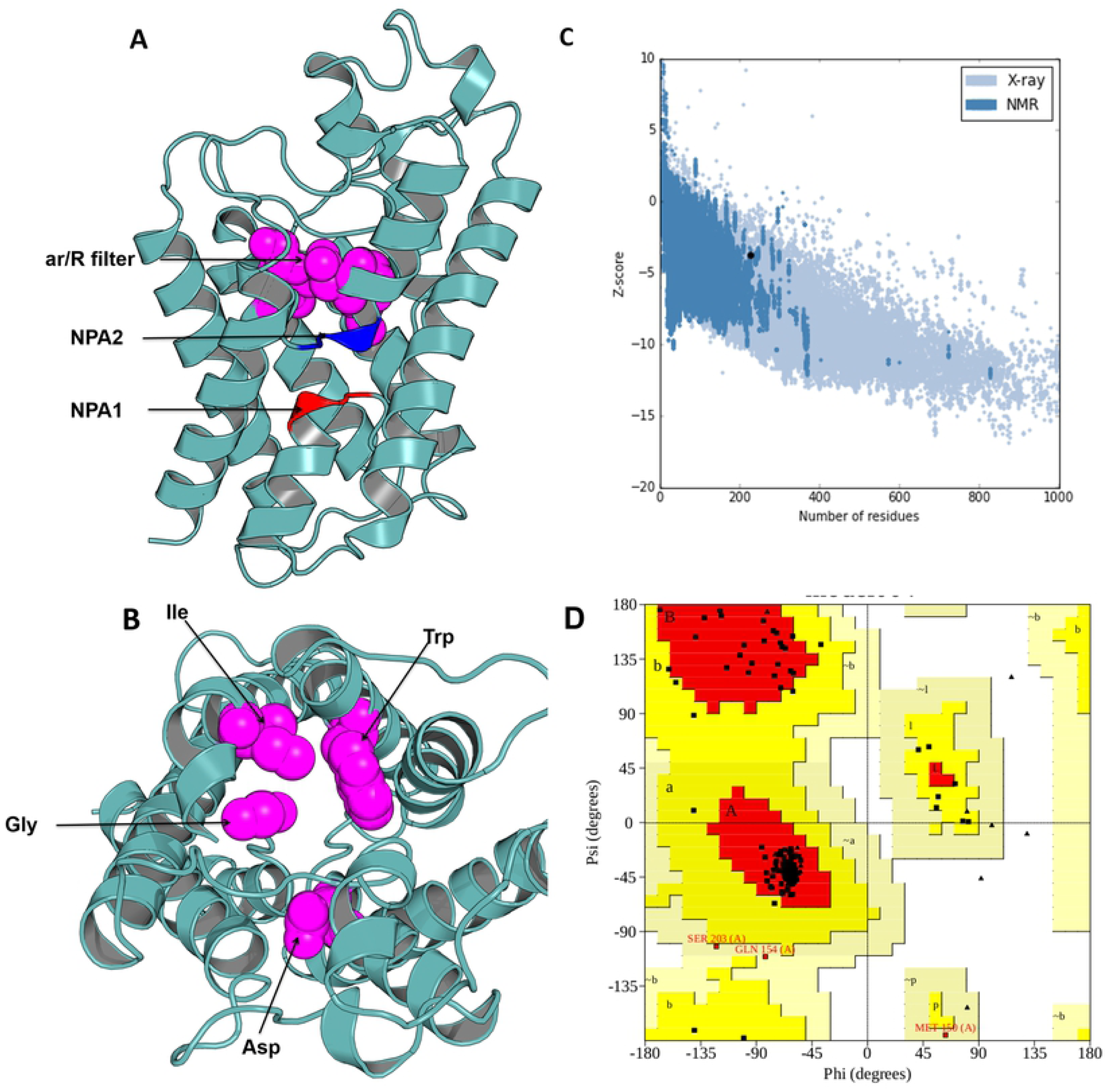
Structural mapping of the AQP1 peptide motifs onto the protein 3D structure. Motifs M7 and M8 were located on the protein surface and in the extra-cellular domain while M9, M10, and M6 were located in the transmembrane domain. The protein was rendered in cartoon (**A)** and protein surface (**B)** formats. The motifs were coloured as follows: M3- red, M7- blue, M8- purple, M9- light green, M10- yellow, and M16- dark green.

**Fig 12.**
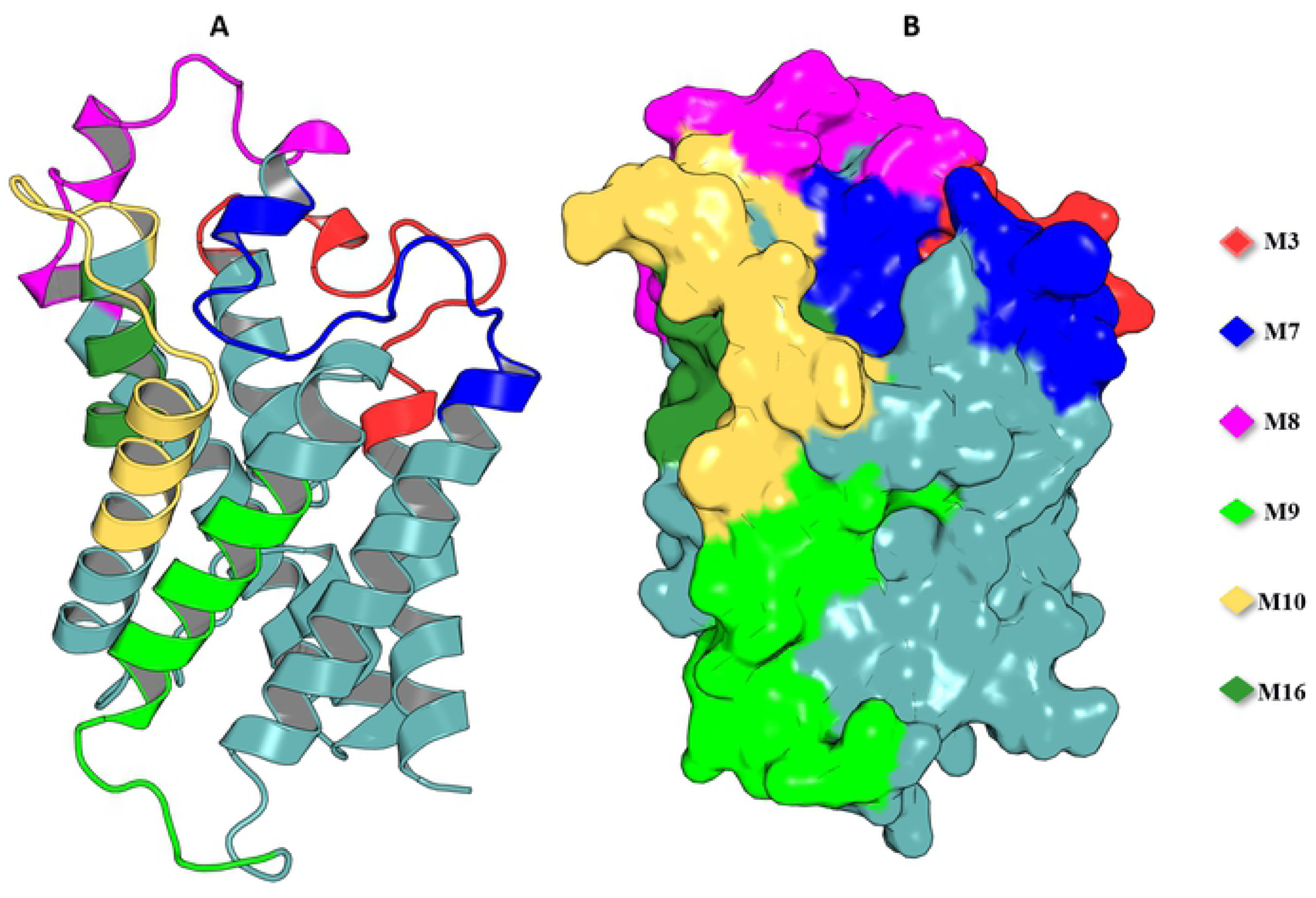
Structural super-imposition of the tick and bovine Aquaporin-1. The protein structures were rendered in cartoon format and the structure super-imposition was done using PYMOL. **A.** The tick AQP1 colored teal, **B.** The bovine 1J4N AQP1 colored purple, and **C.** Cartoons showing superposed structures of tick AQP1 and bovine AQP1

### *In silico* analysis of the peptide motif immunogenicity

Motifs M7 and M8 were the most immunogenic with the highest parker hydrophilicity scores of 1.153 and 2.612 respectively. The immunogenicity of M7 and M8 motifs was further confirmed by the highest BepiPred scores of 0.53 and 0.22 respectively (Table 2).

**Table 2.** Prediction of B cell peptide motif immunogenicity.

| Peptide<br>motif name | Bepipred scores | Chou & Fasman<br>Scores | Karplus and Schulz<br>flexibility | Parker |
| --- | --- | --- | --- | --- |
| M3 | -4.07 | 1.009 | 0.953 | -0.144 |
| M7 | 0.53 | 0.980 | 0.987 | 1.153 |
| M8 | 0.228 | 1.106 | 1.009 | 2.612 |
| M9 | -0.571 | 0.898 | 0.982 | -1.483 |
| M10 | -0.048 | 0.989 | 0.958 | -0.059 |
| M16 | -0.880 | 0.923 | 1.015 | -0.521 |

## Discussion

In this study, *in silico-*based methods were used to predict AQP1 peptide motifs unique to ticks isolated from this study and other AQP1 from the Protein Data Bank. The predicted motifs were further assessed for their potential of inducing humoral immune response and therefore their potential for being good vaccine antigens. We identified a total of 6 unique tick-AQP1 peptide motifs.

Tick AQP1 proteins from this study shown regions of similarity to AQP1 from other tick species and other organisms (Fig 7). This is due to the obvious fact that AQP1 plays similar functions in the species in which it exits. In addition all AQP proteins have 6 transmembrane alpha helical domains, a signature characteristics to all Major Intrinsic Protein (MIP) superfamily proteins [36]. The highest AQP1 sequence similarity within the orthologous proteins was observed mainly around the Asparagine-Proline-Aline (NPA) motif and aromatic/Arginine (ar/R) selectivity filters [38–40]. These filters slow down the flow of molecules across the protein pore by a Grotthuss mechanisms [38–40]. Similar studies have indicated that the arrangement of the ar/R residues directly correlates with the functional properties of the channel [38–40]. Classical water selective aquaporins (CAQPs) such as AQP1 usually show tight ar/R clusters within which passage of water is allowed while ions and glycerol are blocked [36,38,41]. Tick AQP1 from this study contained an aspartic acid residue (D) after the second NPA motif (Fig 7) which is the signature sequence of Aquaglyceroporin (AQGP). This finding is in conformity with previous studies which looked at AQP1 proteins from different tick species [16,41–44]. An aspartic acid residue always coexists with a longer loop which increases the pore’s permeability to larger molecules such as glycerol [45]. Despite the presence of a D residue, this functional characterisation of tick AQP1 does not warrant transportation of glycerol or Urea [13]. Such tight water transport mechanisms seen in tick AQP1 helps them to concentrate the blood meal component in the midgut by removing only water back into the host via the salivary gland [42].

The high sequence similarity within the AQP1 from all different tick species further emphasizes its potential for use as a catch-all anti-tick vaccine candidate [17]. Whole tick AQP1 structure super-imposition to the bovine AQP1 protein indicated that the two proteins were structurally similar (RMS= 1.475). Such similarities with the host protein can affect vaccine efficacy and have potential of causing adverse autoimmune effects if vaccinated with the whole AQP1 recombinant protein [46]. This challenge could have been the reason for failure of the recombinant AQP1 protein as vaccine against *I. ricinus* [15]. Future AQP1-anti-tick vaccines therefore should at least be based on reverse vaccinology methods focusing on unique tick peptide motifs that might constitute vaccine development pipeline [47]. Peptide based vaccines use specific peptide fragments that induce high specific immune response thereby increase vaccine efficacy and reduce potential adverse effects caused by whole protein vaccination [48]. A number of peptide-derived vaccines have been developed, some of these are in clinical trials. A good example is the discovery of neutralizing epitopes to HIV and influenza viruses [49,50].

Multiple sequence alignment of tick AQP1 amino acid sequences and their bovine orthologs highlighted some differences between them. These regions of difference were observed in the loops of both proteins. It is from this region that all tick AQP1 peptide motifs predicted in this study were localized. Structural mapping of peptide motifs predicted in this study onto the modeled 3D aquaporin molecule showed that motifs M9, M16, part of M10 and M3 are all located on tick AQP1 transmembrane domains. Motif modeling and sequence logo identification showed that these motifs contained high number of hydrophobic residues which usually encode for most transmembrane alpha helices [51]. This phenomenon is due to the fact that hydrophobic residues are non-polar thus they have to fold in such a way that they are buried in lipid bilayer of the cell membrane [51].

Moreover, these motifs (M3, M9, M10, and M16) showed low level of immunogenicity and cannot be accessed by antibodies upon cattle vaccination and tick feeding. Only two motifs (M7 and M8) are extracellular AQP1 peptide motifs and are potentially immunogenic and would be easily accessed by antibodies upon cattle vaccination and tick feeding. The average length of all predicted peptide motifs was 20 amino acids, our best two peptides M7 and M8 were each 20 amino acids long (fig 9). Given their good predicted immunogenicity scores, and the fact that they are of appropriate length, they can potentially induce immune response when properly conjugated. Peptides of 8-12 amino acid residues are sufficient enough to be expressed by Major histocompatibility complex (MHCII) to cause an appropriate immune response [52]. These motifs therefore, need to be further evaluated for their immunogenicity using wet-lab animal models.

## Conclusion

We identified two motifs (M7 and M8) which can potentially be incorporated into an AQP1-derived anti-tick vaccine. We also demonstrated high level of amino acid sequence conservation across AQP1 from different tick species. This indicates that if produced, an AQP1-derived anti-tick vaccine might show significant effect against a wide range of tick species. Further studies are needed to test if AQP1-derived motifs can effectively induce a desired immune response and if anti-bodies derived therefrom can effectively prevent tick infestation.

## Acknowledgments

We are grateful to the different veterinary officers in the 4 districts who helped the core study team in collecting tick samples. The authors received no specific funding for this work.

